# GEOexplorer: an R/Bioconductor package for gene expression analysis and visualisation

**DOI:** 10.1101/2021.10.06.459411

**Authors:** Guy P Hunt, Rafael Henkin, Fabrizio Smeraldi, Michael R Barnes

## Abstract

**Background:** Over the past three decades there have been numerous molecular biology developments that have led to an explosion in the number of gene expression studies being performed. Many of these gene expression studies publish their data to the public database GEO, making them freely available. By analysing gene expression datasets, researchers can identify genes that are differentially expressed between two groups. This can provide insights that lead to the development of new tests and treatments for diseases. Despite the wide availability of gene expression datasets, analysing them is difficult for several reasons. These reasons include the fact that most methods for performing gene expression analysis require programming proficiency.

**Results:** We developed the GEOexplorer software package to overcome several of the difficulties in performing gene expression analysis. GEOexplorer was therefore developed as a web application, that can perform interactive and reproducible microarray gene expression analysis, while producing a wealth of interactive visualisations to facilitate result exploration. GEOexplorer is implemented in R using the Shiny framework and is fully integrated with the existing core structures of the Bioconductor project. Users can perform the essential steps of exploratory data analysis and differential gene expression analysis intuitively and generate a broad spectrum of publication ready outputs.

**Conclusion:** GEOexplorer is distributed as an R package in the Bioconductor project (http://bioconductor.org/packages/GEOexplorer/). GEOexplorer provides a solution for performing interactive and reproducible analyses of microarray gene expression data, empowering life scientists to perform exploratory data analysis and differential gene expression analysis on GEO microarray datasets.

## Background

Over the past three decades, there have been many gene expression developments, including the creation of microarray techniques, that have led to an explosion in the number of gene expression studies being performed. Many of these studies publish their datasets in the Gene Expression Omnibus (GEO), which is a public repository that makes gene expression datasets freely available to the public [1].

Gene expression analysis is used in a broad range of research areas including cancer research such as breast cancer [2] and prostate cancer [3], understanding current epidemics and pandemics for instance obesity [4] and COVID-19 [5], and drug discovery such as treating inflammatory bowel disease using topiramate and treating lung carcinomas using cimetidine [6]. Gene expression analysis has been used to uncover new insights in disease pathology as well as identify targets for drugs and diagnostic tests. Gene expression analysis usually occurs in two stages. The first stage is called “Exploratory Data Analysis” (EDA) which is used to gain an overall understanding of the gene expression dataset. The second stage is called “Differential Gene Expression Analysis” (DGEA) which identifies the genes that are expressed differently between two groups.

Despite the availability of microarray gene expression datasets, performing analysis on the datasets can be difficult. The reasons for that include the high dimensionality of the datasets, variation in experiment structures and the statistical knowledge required to perform the analysis. There are several R packages developed to overcome these challenges. However, the use of the packages requires programming proficiency. As many life scientists are not proficient at programming, this represents a hurdle to performing gene expression analysis. The website GEO2R, https://www.ncbi.nlm.nih.gov/geo/geo2r/, enables users to perform gene expression analysis, on microarray datasets, without requiring programming proficiency [1]. GEO2R substantially meets the need for an intuitive and accessible interface to perform gene expression analysis on microarray datasets.

GEOexplorer aims to expand upon the foundations laid by GEO2R and overcome several additional challenges to performing gene expression analysis. Specifically, GEOexplorer aims to enable users with no programming skills to perform gene expression analysis, provide a rich selection of analytical techniques and publication ready figures, and enable in-depth exploration of the gene expression analysis results, using interactive visualisations.

Based on the feedback from life scientists, we developed GEOexplorer as an interactive Shiny [7] web-based application in the GEOexplorer R/Bioconductor package. GEOexplorer provides an integrated platform for extracting, transforming, analysing, visualising, and exploring GEO microarray datasets.

GEOexplorer takes as input a GEO series ID, also known as a GEO accession code, which it uses to extract the microarray gene expression dataset and experimental information from GEO. GEOexplorer makes performing EDA and DGEA easy by requiring minimal configurations to be set, allowing the analysis to be automated. GEOexplorer also adheres to the FAIR Guiding Principles for scientific data [8]. GEOexplorer delivers a wide range of information-rich visualisations and tables, for both EDA and DGEA, which form a comprehensive, transparent, and reproducible analysis of microarray data.

GEOexplorer is available at http://bioconductor.org/packages/GEOexplorer/. The application can be deployed as a standalone web-service, as we did for the publicly hosted version available at https://geoexplorer.shinyapps.io/GEOexplorer, where the full functionality of the application can be explored.

## Implementation

### General design of GEOexplorer

GEOexplorer is written in the R programming language. GEOexplorer connects the functionality of several widely used packages available from Bioconductor and CRAN. As a result, GEOexplorer can deliver multiple types of outputs and visualisations to easily translate microarray datasets into knowledge and insights. GEOexplorer uses the *limma* package to perform DGEA as it was found to be among the best performing packages for microarray datasets [9].

The web application and all its features are provided by a call to the *loadApp* function, which fully exploits the Shiny reactive programming paradigm to efficiently (re-)generate the rendered components and outputs upon detection of changes in the input widgets.

The layout of the user interface is built with a sidebar containing widgets for the data collection and transformation options. The main panel is structured with three different tabs that mirror the different steps to perform gene expression analysis. The steps include reviewing: the dataset information, the results of EDA, and the results of DGEA. The widgets for DGEA options are available within the DGEA tab. Each tab has multiple subtabs to guide the user through a logical workflow. Additionally, GEOexplorer contains features like tooltips, based on the bootstrap components in the *shinyBS* package [10], to guide users through various steps.

The *plotly* [11] graphics system is used to generate interactive visualisations, enabling interactions by brushing or clicking on them in the Shiny framework. Interactive heatmaps are generated with the *heatmaply* package [12], and tables are displayed as interactive objects for efficient navigation via the *DT* package [13].

The functionality of GEOexplorer is extensively described in the package vignette and is generated via the Bioconductor build system. Documentation for each function is provided, with examples that can be seen on the vignette at http://bioconductor.org/packages/devel/bioc/vignettes/GEOexplorer/inst/doc/GEOexplorer.html. Additionally, a YouTube video tutorial demonstrating GEOexplorer’s functionality is available at https://youtu.be/8R8yqMlPCVM.

### System requirements

GEOexplorer has been tested on macOS, Linux, and Windows. It can be downloaded from the Bioconductor project page (http://bioconductor.org/packages/GEOexplorer/), and its development version can be found at https://github.com/guypwhunt/GEOexplorer/.

Since GEOexplorer is normally installed on local systems, its speed and performance will vary depending on the hardware specifications available. In our experience, a typical modern laptop or workstation with at least 8 GB RAM is sufficient for GEOexplorer to process nearly all GEO microarray datasets. However, a modern laptop or workstation with as little as 1 GB RAM was sufficient to process most GEO microarray datasets. The size of the dataset, data quality and the number of experimental conditions included in DGEA are the main factors that influence the time required to complete a session with GEOexplorer, after familiarizing with its interface and workflow etc.

### Typical usage workflow

During the typical usage session of GEOexplorer (Fig. 1), users need to input a GEO accession code. The GEO accession code can be identified from the National Center for Biotechnology Information GEO, https://www.ncbi.nlm.nih.gov/geo/. GEOexplorer then sources the microarray gene expression dataset and experimental study information from GEO. The user can then select the platform of interest and transformation options. GEOexplorer then displays information about the study and experiment as well as performing and displaying the results of EDA. GEOexplorer forces users to perform EDA on the GEO microarray dataset, as this is a fundamental requirement for setting the configurations for DGEA. The DGEA options can then be selected based on the outputs of EDA. Finally, the outputs of DGEA can be reviewed.

**Figure 1:**
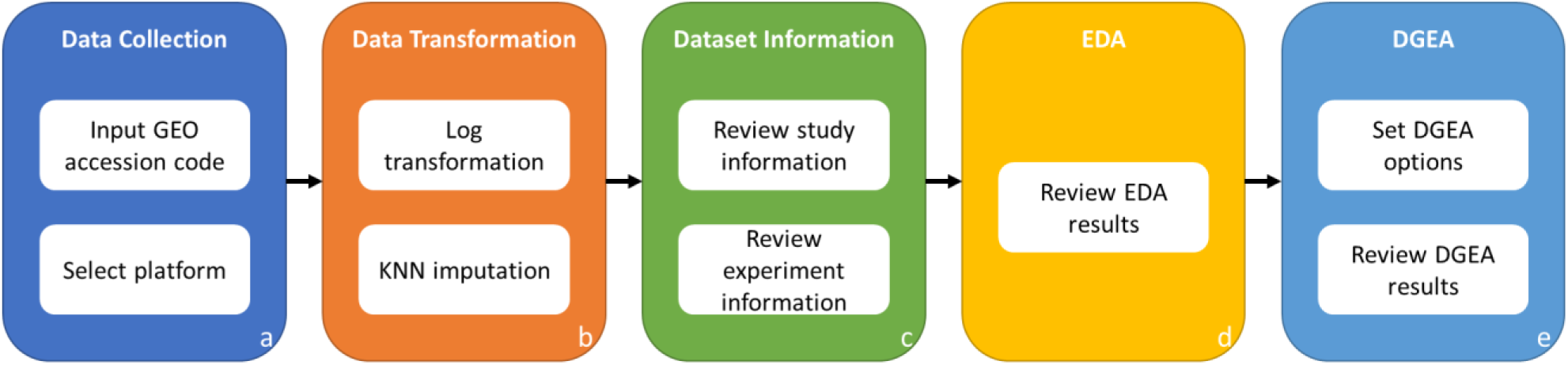
GEOexplorer workflow overview. **a** gene expression analysis with GEOexplorer starts by inputting the GEO accession code of the interested microarray study. The platform of interest is then selected, each platform is a microarray experiment that was performed as part of the study. **b** log transformation and k-nearest neighbour (KNN) imputation can be selected before analysing the data. **c** the dataset information including information about the study and experiment can be reviewed. **d** the results of exploratory data analysis can be reviewed. **e** the options for differential gene expression analysis can be set based on the outputs of exploratory data analysis. Subsequently, the outputs of differential gene expression analysis can be reviewed.

For each setting in the main application, GEOexplorer gives a text description of the setting and its effect. All figures generated and displayed in the user interface can also be downloaded with a simple mouse click. Additionally, users can explore each of the interactive figures.

## Results

The functionality of GEOexplorer is described in the next sections and is illustrated in detail for the analysis of space-flown mice thymus microarray data (published in [14]).

### Data collection and transformation

The setup for the data collection and transformation is carried out in the sidebar panel (Fig. 2). The user needs to input the GEO accession code into GEOexplorer. GEOexplorer then sources the GEO file, containing the microarray gene expression dataset(s) and study information, directly from GEO and parses it using the *GEOquery* package [15]. Once the GEO file has been downloaded and parsed, the user can then configure which platform to use. Each platform relates to a different microarray gene expression dataset from the study.

**Figure 2:**
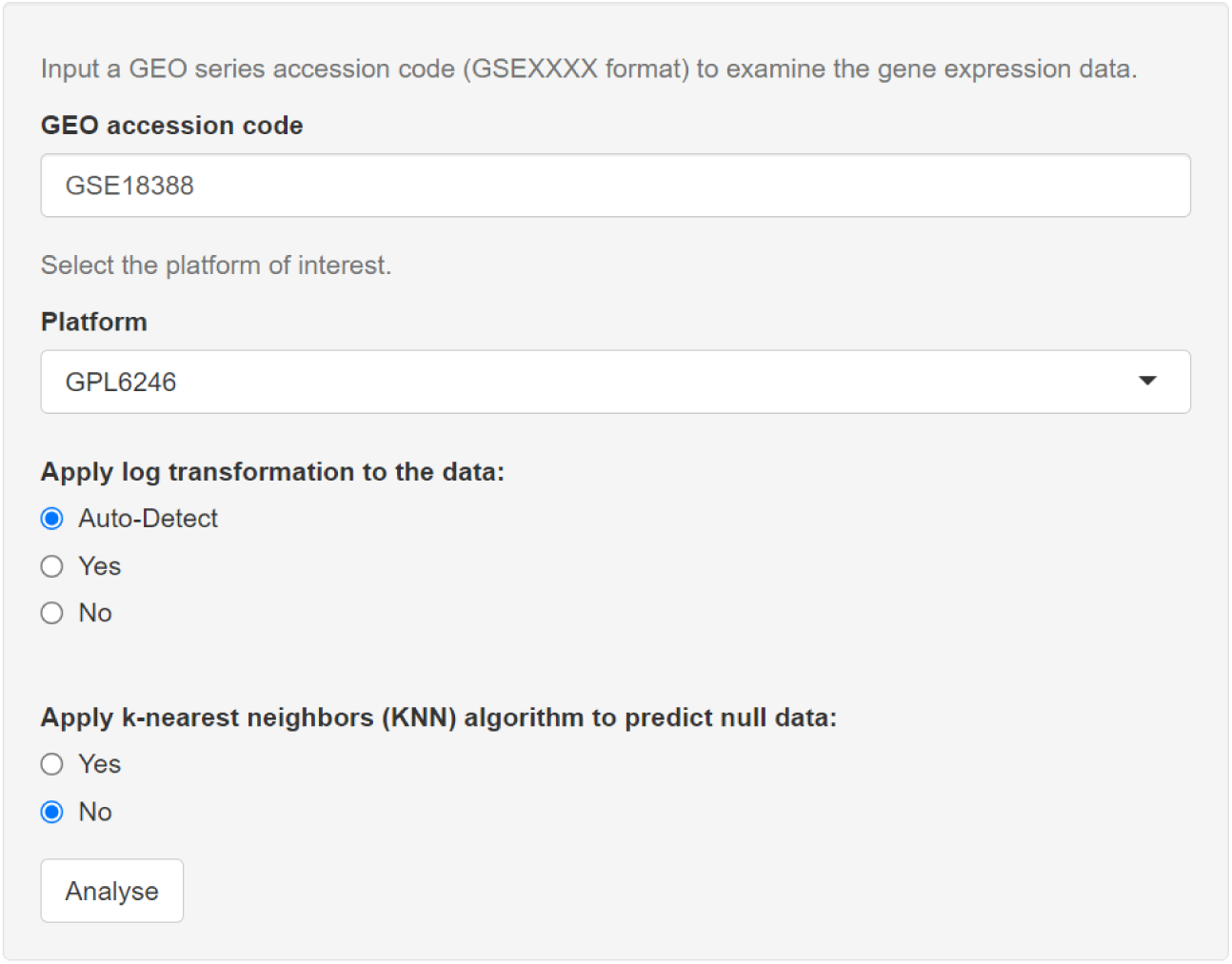
GEOexplorer data sourcing and transformation settings.

The user can select whether to apply log transformation to the expression data and whether to use KNN imputation to fill in missing values within the expression data. KNN is applied using the *impute* package [16]. By default, GEOexplorer will automatically identify if log transformation should be applied to the expression dataset.

### Study and experiment information

After the user clicks the analyse button in the sidebar, information about the study and experiment is displayed in the *Dataset Information* tab. Specifically, an overview of the study is available in the *Experiment Information* subtab (Fig. 3a), the experiment conditions are available in the *Experimental Conditions Information* subtab (Fig. 3b) and the gene expression dataset is available in the *Gene Expression Dataset* subtab (Fig. 3c). The gene expression dataset is displayed post log transformation and KNN imputation, if selected, and therefore is used to see the effects of log transformation and KNN imputation on the gene expression dataset.

**Figure 3:**
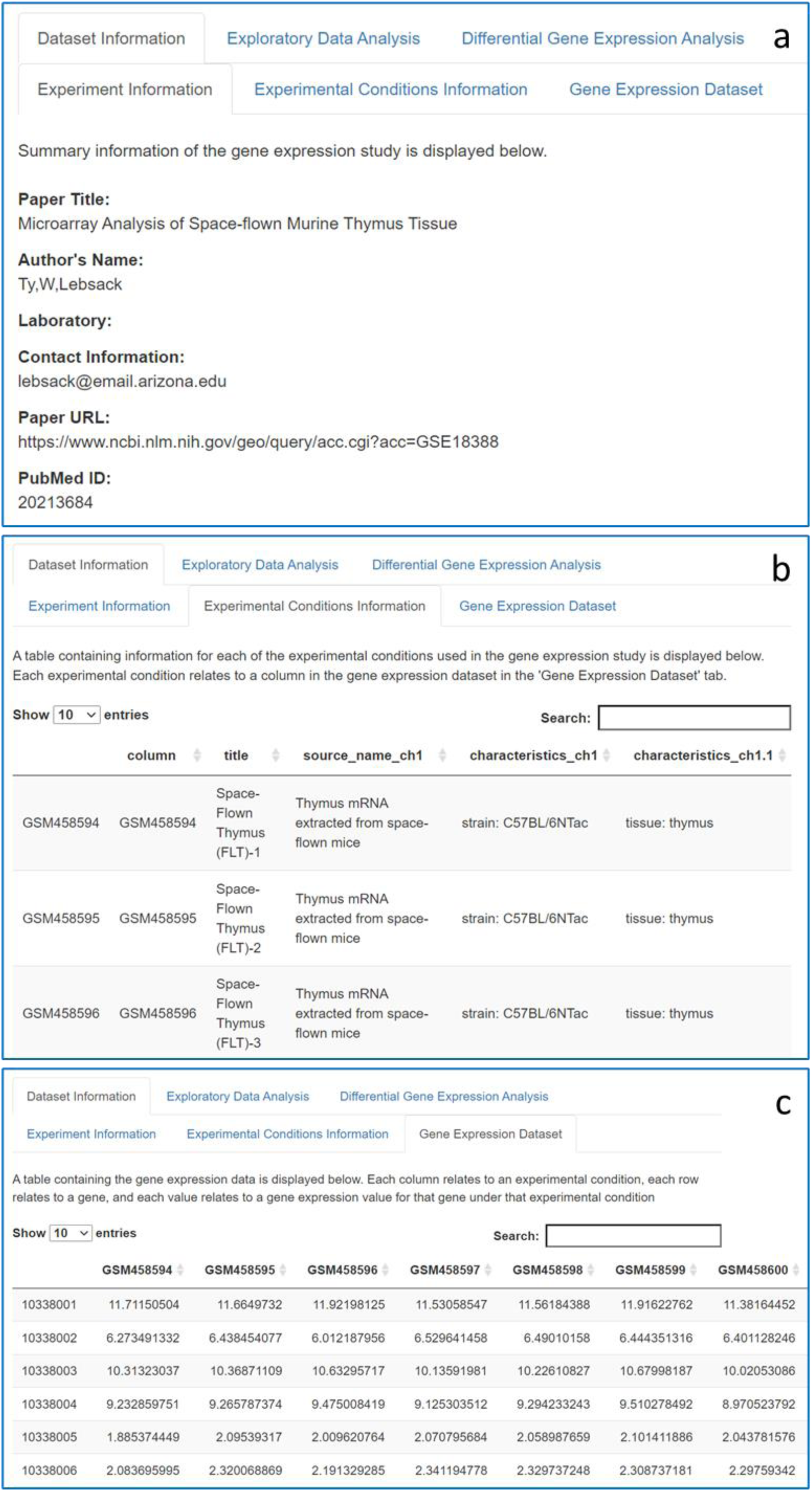
Screenshots of the GEOexplorer study and experiment information. **a** Experiment information. **b** Experimental conditions information. **c** Gene expression dataset.

### Generating and exploring the results for EDA

The expression dataset post data transformation is used as the basis of EDA. It, therefore, incorporates the log transformation and KNN configurations set during data transformation.

GEOexplorer automatically performs several EDA techniques and displays the results in interactive visualisations after the user clicks the analyse button in the sidebar (Fig. 2). Each of the visualisations can be downloaded from GEOexplorer and used in publications.

GEOexplorer displays an interactive box-and-whisker plot (Fig. 4a). The box-and-whisker plot displays the distribution of gene expression values for each experimental condition including the min, max, median, 1^st^ quartile and 3^rd^ quartile. The box-and-whisker plot is useful for identifying if the experimental conditions’ gene expression values are median-centred. Median-centred values indicate the gene expression data is normalised and therefore comparable during DGEA. If the experimental conditions are not median-centred, then the user can configure forced normalisation during DGEA.

**Figure 4:**
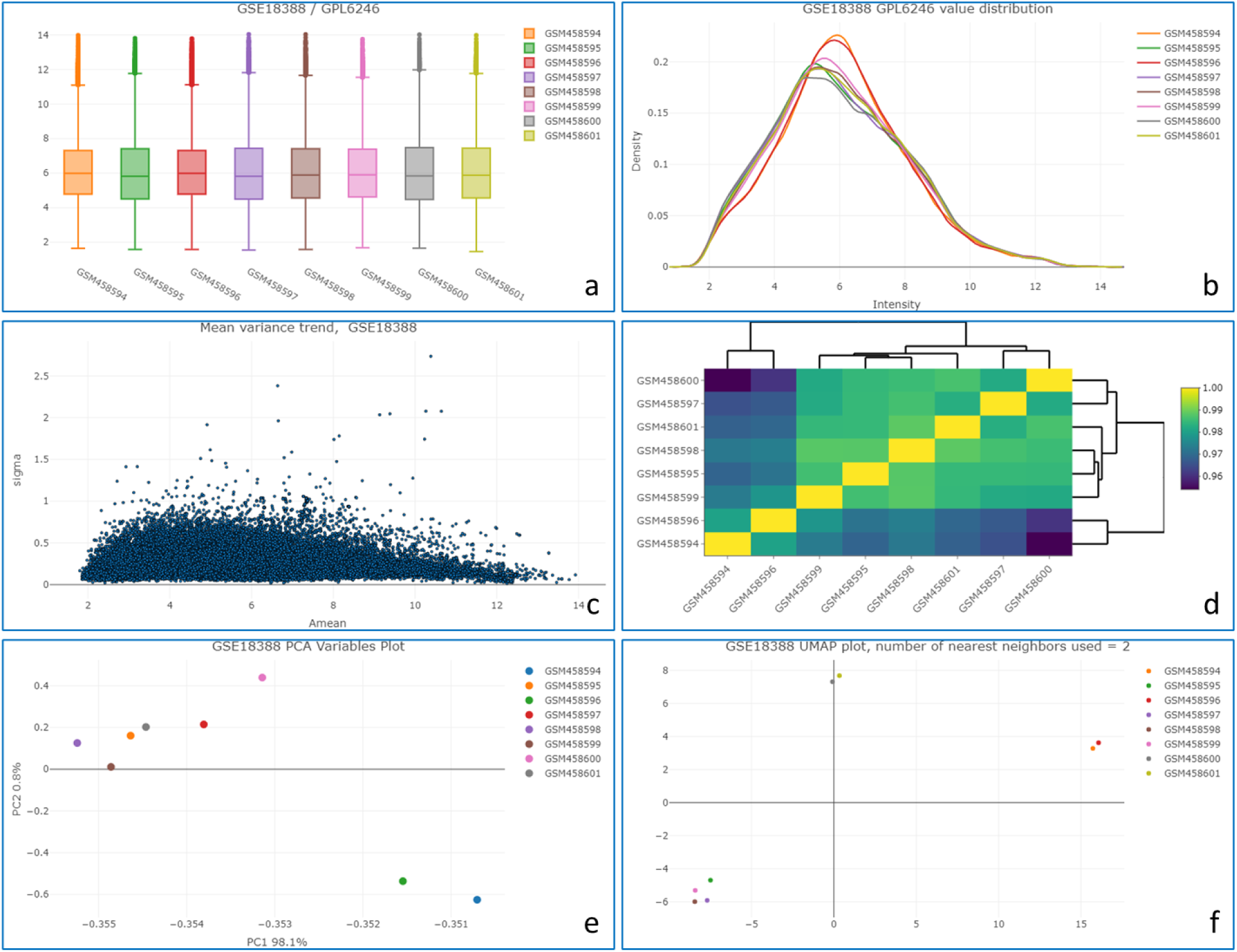
Screenshot selection of the GEOexplorer Exploratory Data Analysis outputs. **a** Box-and-whisper plot. **b** Expression density plot. **c** Mean-variance plot. **d** Heatmap plot. **e** Principal component analysis (PCA) variables plot. **f** Uniform manifold approximation and projection (UMAP) plot.

GEOexplorer displays two interactive expression density plots, one which is 2D (Fig. 4b) and one which is 3D. Firstly the density distributions of each experimental condition are calculated using the *stats* R package. The density distributions of each experimental condition are then plotted on an interactive scatter graph. Like the box-and-whisker plot, the expression density plots are useful for identifying if the expression data needs to be normalised for DGEA. If the density curves do not appear to be normally distributed, then it is advisable to force normalisation during DGEA.

GEOexplorer displays an interactive mean-variance plot (Fig. 4c). The mean-variance plot displays the log residual standard deviation versus the average log expression of the linear model for each gene. This can be used to assess if there is a lot of variation within the gene expression dataset after fitting it to a linear model. The level of variation can be used to determine whether to apply the precision weights option during DGEA. If there is a strong mean-variance trend in the gene expression dataset the precision weights can improve the accuracy of DGEA. The linear model fitting is performed using the *limma* R package [9].

GEOexplorer displays an interactive heatmap (Fig. 4d). The covariances/correlations between each of the experimental conditions are calculated and then displayed in a heatmap. The heatmap is useful for identifying experimental conditions that have closely related gene expression values. The covariances/correlations are calculated using the *stats* R package. The covariances/correlations are then plotted on an interactive heatmap using the *heatmaply* R package [12].

GEOexplorer performs PCA and displays the results in three interactive plots. GEOexplorer uses the *prcomp* function from the *stats* R package to perform PCA.

The first PCA plot is a scree plot. The scree plot displays the percentage of the variance in the gene expression dataset captured by each principal component.

The second PCA plot is an interactive individuals plot. This is useful for identifying genes with similar variations across the experimental conditions. The principal component values for the genes are extracted using the *factoextra* R package [17]. For each gene, the value of the principal component with the greatest variance is plotted against the value for the principal component with the second greatest variance.

The final PCA visualisation is an interactive variables plot (Fig. 4e). Like the heatmap plot, the variables plot is useful for identifying similar experimental conditions that could be grouped during DGEA. The principal component values for the experimental conditions are extracted using the *factoextra* R package [17]. For each experimental condition, the value of the principal component with the greatest variance is plotted against the value for the principal component with the second greatest variance.

GEOexplorer displays an interactive UMAP plot (Fig. 4f) [18]. Like the heatmap and variables plot, the UMAP plot helps to group similar experimental conditions. These groups can then be used in DGEA. The number of KNN used can be updated by the user. Each experimental condition in the dataset is given an edge to each of its KNNs. This helps to group experimental conditions with their nearest neighbours. The *umap* R package [19] is used to reduce the dimensionality of the gene expression dataset to two dimensions.

### Generating and exploring the results for DGEA

The expression dataset post data transformation is used as the basis of DGEA. It, therefore, incorporates the log transformation and KNN configurations set during data transformation.

To perform DGEA the user must select the experimental conditions to include in group 1 and group 2 (Fig. 5a). The gene expression values for the experimental conditions in group 1 will then be compared to the gene expression values for the experimental conditions in group 2. To estimate the variability and perform the statistical tests required to perform DGEA, 1 group needs at least 2 experimental conditions and the other group needs at least 1 experimental condition. Therefore, if the microarray gene expression dataset has less than three experimental conditions, DGEA cannot be performed.

**Figure 5:**
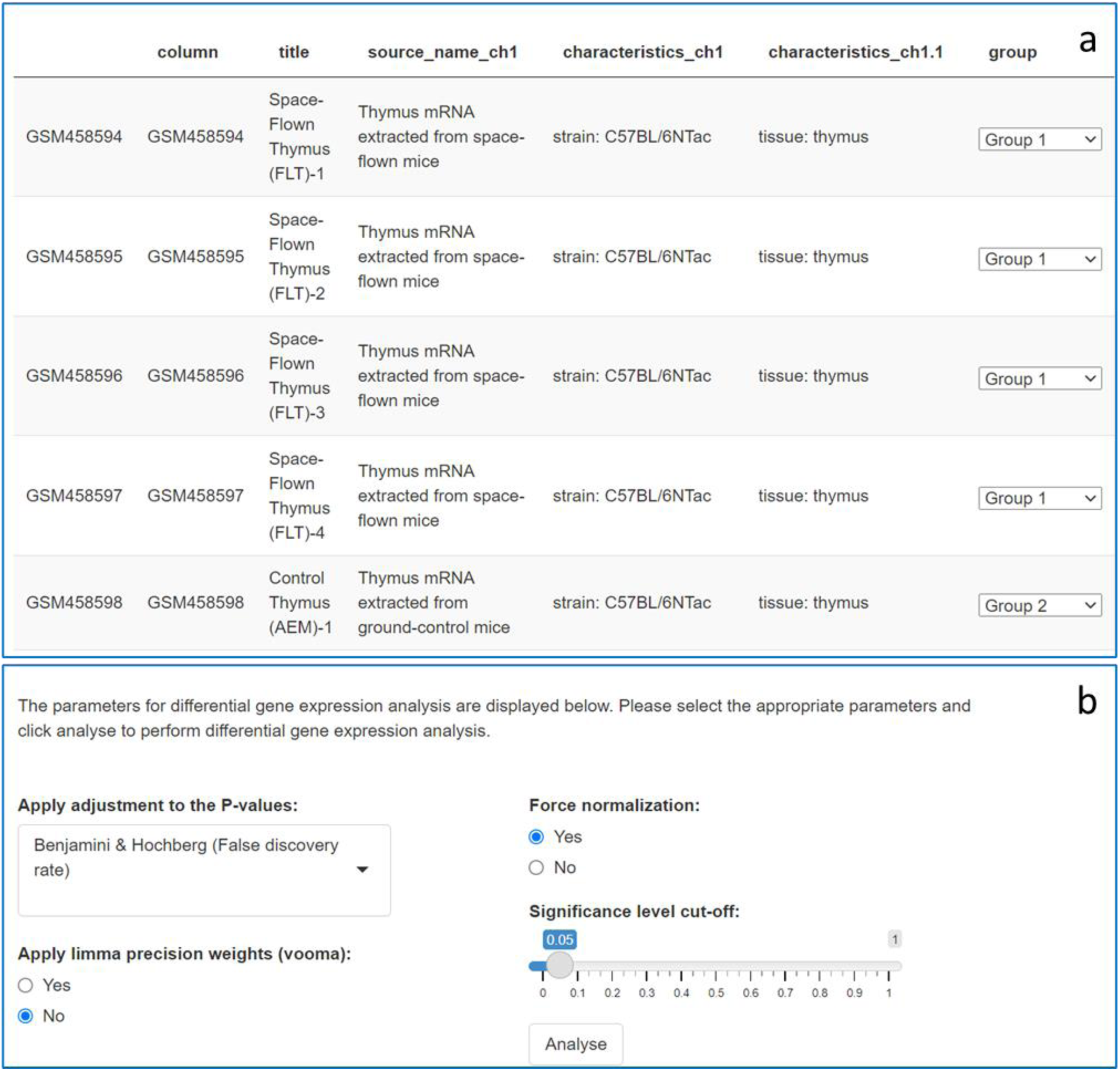
GEOexplorer differential gene expression analysis settings.

Several different options can be selected during DGEA (Fig. 5b). This includes selecting from the following P-value adjustments; Benjamini & Hochberg (False discovery rate), Benjamini & Yekutieli, Bonferroni, Holm, or none. Whether to use limma precision weights, to force normalisation and the significance level cut off can all be selected. After selecting the options, clicking the analyse button triggers DGEA. GEOexplorer uses multiple functions from the *limma* R package [9] to perform DGEA.

If the user selected forced normalisation, the expression intensities across the two groups of experimental conditions are normalised and therefore consistent.

If the user selects to use limma precision weights, then the mean-variance relationship is estimated, and the observational-level weights are calculated from the mean-variance relationship.

DGEA is done by fitting a linear model to each gene within the dataset. The linear model estimates the fold change in the gene’s expression while considering standard errors by applying empirical Bayes smoothing. Genes are subsequently ranked based upon their fold change values.

GEOexplorer displays the results of DGEA in several visualisations (Fig. 6) to help explore the results. These include a table of the top differentially expressed genes, a histogram, a Venn diagram, a QQ plot, a volcano plot, and a mean difference plot. Each of these visualisations can be downloaded from GEOexplorer to be used in publications.

**Figure 6:**
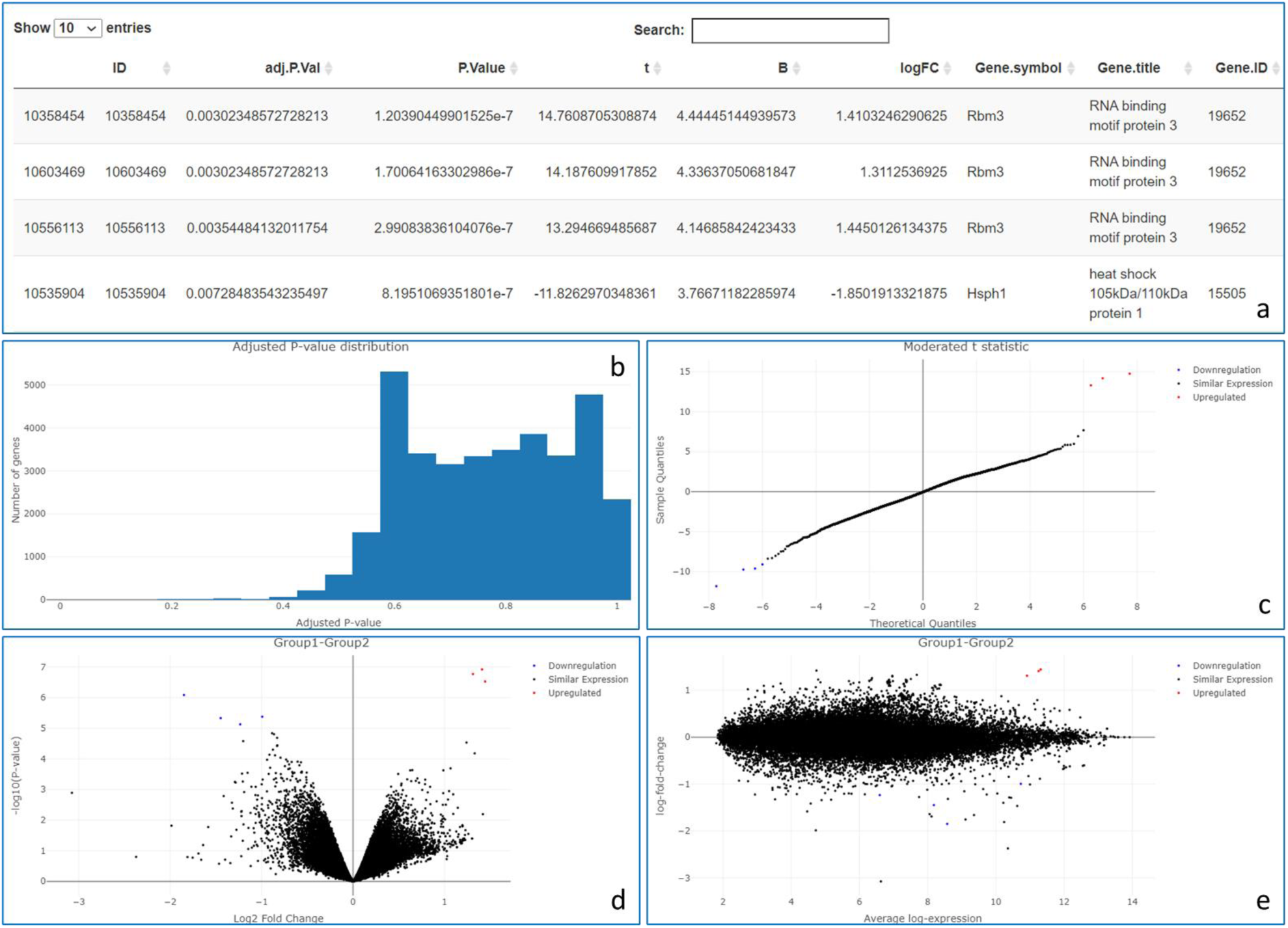
Screenshot selection of the GEOexplorer differential gene expression analysis outputs. **a** Table of top 250 differentially expressed genes. **b** Histogram plot of adjusted P-values. **c** Quantile-quantile (QQ) plot. **d** Volcano plot. **e** Mean-difference plot.

The table of the top differentially expressed genes contains the 250 most differentially expressed genes (Fig. 6a). The table incorporates the P-value adjustment selected by the user.

The interactive histogram displays the distribution of adjusted P-values across all the genes as per (Fig. 6b). This is useful for determining if an appropriate P-value adjustment was selected.

Based on the adjusted P-value each gene is identified as upregulated, downregulated, or not significantly expressed differently in group 1 compared to group 2.

The Venn diagram displays the set of the genes that are either upregulated or downregulated and the set of the genes that are not significantly expressed differently between the two groups of experimental conditions. Like the histogram plot, the Venn diagram plot is useful for determining if an appropriate P-value adjustment was selected.

The QQ plot displays the quantiles of the differentially expressed genes plotted against the theoretical quantiles of a Student’s t distribution for each gene (Fig. 6c).

The volcano plot displays the statistical significance (-log10 P-value) versus magnitude of change (log2 fold change) for each gene (Fig. 6d).

The mean difference plot displays the log2 fold change against the average log2 expression values for each gene (Fig. 6e).

In the QQ, volcano and mean difference plots, the upregulated genes are highlighted in red, and the downregulated genes are highlighted in blue.

## Discussion

The principal aims of GEOexplorer were to enable life scientists to perform gene expression analysis on GEO microarray datasets without the need to be proficient at programming, provide a wide selection of analytical techniques and output the results as interactive visualisations to make exploring the results easy.

GEOexplorer integrates several R packages from Bioconductor and CRAN and as a result, incorporates several of the best R packages for performing gene expression analysis on GEO microarray datasets. GEOexplorer is itself part of Bioconductor, and thus is integrated into a build system that continuously checks all its components and their interoperability. This guarantees that GEOexplorer’s features correctly interface with the latest version of its package dependencies. Notably, the Bioconductor project enforces several best practices to enhance the usability of its components, with both internal and external documentation (for the individual functions and as complete tutorials, in the form of vignettes), as well as providing unit test sets to ensure the software is working as expected. GEOexplorer adheres to these guidelines, which can be essential to define robust software [20] that can be adopted by a wide range of users.

The rich selection of analysis and interactive outputs constitutes a significant advantage for uncovering new insights from GEO microarray datasets. GEOexplorer provides a platform to facilitate discoveries in a standardized way, which consequently improves the reproducibility of the analyses.

## Conclusion

The infrastructure provided by the GEOexplorer R/Bioconductor package delivers a web browser application that guarantees ease of use through interactivity, together with reproducible research, for the essential steps of EDA and DGEA investigation in microarray analysis. The combination of these features is a key factor for efficient, quick, and robust extraction of information, while leveraging the facilities available in the Bioconductor project in terms of analytical and statistical methods.

The wealth of information users can extract using GEOexplorer, through its rich selection of analysis and interactivity, may play a critical role when individuals choose a tool to adopt in their projects. Furthermore, the way GEOexplorer guides life scientists through EDA and DGEA reduces the amount of bioinformatics/biostatistics expertise required to perform and understand gene expression analysis. The design choices for GEOexplorer aim at making gene expression analysis as simple as possible.

Our package reaches out to life scientists, being simple to install and use, without the need to be proficient at programming, based on robust statistical methods, and offering multiple levels of documentation. GEOexplorer enables life scientists to easily take control of the analysis of GEO microarray datasets, while providing an accessible framework for reproducible analysis.

## Additional information

### Availability and requirements

**Project name:** GEOexplorer

**Project home page:** http://bioconductor.org/packages/GEOexplorer/ (release) and https://github.com/guypwhunt/GEOexplorer (development version)

**Archived version:** https://doi.org/10.5281/zenodo.5511937, package source as a gzipped tar archive of the version reported in this article

**Project documentation:** rendered at http://bioconductor.org/packages/GEOexplorer/

**Operating systems:** Linux, Mac, Windows

**Programming language:** R

**Other requirements:** R 4.0 or higher and Bioconductor 3.14 or higher

**License:** GPL 3.0 or higher

**Any restrictions to use by non-academics:** none

## Abbreviations

DGEA: Differential Gene Expression Analysis
EDA: Exploratory Data Analysis
GEO: Gene Expression Omnibus
KNN: K-Nearest Neighbour
PCA: Principal Component Analysis
QQ: Quantile-Quantile
UMAP: Uniform Manifold Approximation and Projection

## Acknowledgements

We thank GEO2R for their excellent code which guided the development of GEOexplorer.

## Funding

Not applicable.

## Availability of data and materials

Data used in the described use cases is available from the following article:

- The space-flown mice thymus microarray is included in PubMed ID: 20213684 (https://doi.org/10.1002/jcb.22547). GEO entry: GSE18388.

The GEOexplorer package can be downloaded from its Bioconductor page http://bioconductor.org/packages/GEOexplorer/ or the GitHub development page https://github.com/guypwhunt/GEOexplorer/.

## Authors’ contributions

G.H. - Methodology, Software Implementation, Writing - Original Draft Preparation, Writing - Review & Editing

R.H, F.S. and M.B – Conceptualisation, Supervision, Writing - Review & Editing

All authors read and approved the final version of the manuscript.

## Ethics approval and consent to participate

Not applicable.

## Consent for publication

Not applicable.

## Competing interests

The authors declare that they have no competing interests.

## Notes

### Competing Interest Statement

The authors have declared no competing interest.

